# Interhemispheric Integration for Complex Behaviors, Absent the Corpus Callosum in Normal Ontogeny

**DOI:** 10.1101/271072

**Authors:** Elliot A. Layden, Kathryn E. Schertz, Sarah E. London, Marc G. Berman

**Affiliations:** Department of Psychology, The University of Chicago, Chicago, IL 60637.; Grossman Institute for Neuroscience, Quantitative Biology and Human Behavior, The University of Chicago, Chicago, IL 60637; The Institute for Mind and Biology, The University of Chicago, Chicago, IL 60637

**Keywords:** functional homotopy, resting-state fMRI, zebra finch, corpus callosum, interhemispheric coordination

## Abstract

Functional homotopy, or synchronous spontaneous activity between symmetric, contralateral brain regions, is a fundamental characteristic of the mammalian brain’s functional architecture(1–6). In mammals, functional homotopy may be predominantly mediated by the corpus callosum (CC), a white matter structure thought to balance the interhemispheric coordination and hemispheric specialization critical for many complex brain functions, including lateralized human language abilities(7, 8). The CC first emerged with the Eutherian (placental) mammals ~160 MYA and is not found in other vertebrates(9, 10). Despite this, other vertebrates also exhibit complex brain functions requiring bilateral integration and lateralization(11). For example, much as humans acquire speech, the zebra finch (*Taeniopygia guttata*) songbird learns to sing from tutors and must balance hemispheric specialization(12) with interhemispheric coordination to successfully learn and produce song(13). We therefore tested whether the zebra finch brain also exhibits functional homotopy despite lacking the CC. Implementing custom resting-state fMRI (rs-fMRI) functional connectivity (FC) analyses, we demonstrate widespread functional homotopy between pairs of contralateral brain regions required for learned song but which lack direct anatomical projections (i.e., structural connectivity; SC). We believe this is the first demonstration of functional homotopy in a non-Eutherian vertebrate; however, it is unlikely to be the only instance of it. The remarkable congruence between functional homotopy in the zebra finch and Eutherian brains indicates that alternative mechanisms must exist for balanced interhemispheric coordination in the absence of a CC. This insight may have broad implications for understanding complex, bilateral neural processing across phylogeny and how information is integrated between hemispheres.

**Significance Statement:** The mammalian brain exhibits strongly synchronized hemodynamic activity (i.e., functional connectivity) between symmetric, contralateral (i.e., homotopic) brain regions. This pattern is thought to be largely mediated by the corpus callosum (CC), a large white matter tract unique to mammals, which balances interhemispheric coordination and lateralization. Many complex brain functions, including human language, are thought to critically rely upon this balance. Despite lacking the CC, the zebra finch exhibits a song learning process with striking parallels to human speech acquisition, including lateralization and interhemispheric coordination. Using resting-state fMRI, we show that the zebra finch brain exhibits widespread homotopic functional connectivity within a network critical for learned song, suggesting that this symmetrical activity pattern may phylogenetically precede the evolution of the CC.

---

Functional homotopy is consistently identified across Eutherian mammals, including humans(3–6), macaques(3), mice(1), and rats(2), using measurement techniques such as rs-fMRI FC(3–6), local field potentials(2), and calcium imaging(1). Homotopic functional connections are among the strongest in the Eutherian brain(3–5), and they are known to shape the network dynamics of non-homotopic connections(5). Additionally, functional homotopy is clinically relevant, as reductions in homotopy are implicated in several human neuropsychiatric disorders(5).

Convergent evidence indicates that the CC is an important mediator of functional homotopy. Functional homotopy is acutely diminished in humans following complete section of the CC(14) and the signal conduction efficacy of intact CC fibers is positively associated with functional homotopy(3, 15). Despite this, humans born without a CC have been found to exhibit normal functional homotopy by adulthood(16), indicating that alternative, compensatory plasticity permits the establishment of functional homotopy(17). This finding suggests that functional homotopy can utilize alternative mechanisms to the CC and supports our postulation that organisms which lack a CC may also exhibit functional homotopy.

In humans, language learning is one of the most complex multi-modal distributed brain processes(18). Human language processing is strongly lateralized, enabling more efficient processing of specialized language tasks(19). Notably greater lateralization for language is associated with reductions in functional homotopy(8). Like humans, zebra finches exhibit vocal learning for song, a process that exhibits deep genomic, neural, behavioral, developmental, and social parallels with human speech acquisition(20). Similar to humans, zebra finches exhibit lateralization of vocal learning(12) and require interhemispheric coordination of premotor activity to enable vocal production(13).

Despite these parallels, the zebra finch brain is not thought to exhibit direct interhemispheric neural projections connecting the homotopic regions of the telencephalon considered essential for learned song(13). The evolutionarily ancient anterior commissure, which comprises the largest interhemispheric projection in the avian brain, connects bilateral regions of caudal telencephalon(21). However, in contrast to the mammalian CC, its projections are primarily heterotopically organized and unidirectional, with the exception of a relatively small arcopallial and amygdaloid cluster (13). While the precise role (i.e., excitation vs. inhibition) of the CC remains controversial(7), the mammalian CC provides a clear mechanism for the maintenance of the hemispheric lateralization and the interhemispheric coordination needed for human language processing. It is less clear, however, how the zebra finch brain maintains this balance without a CC.

Our primary hypothesis was that, if interhemispheric coordination is necessary to support vocal learning and production, the zebra finch should exhibit a pattern of functional homotopy analogous to Eutherians, despite lacking the CC. Second, given that only male zebra finches sing, and males exhibit sex-specific structural(22) and functional(12) lateralization within the brain network for learned song, we predicted that males would exhibit less functional homotopy than females within this network, directly parallel to the association between language lateralization and functional homotopy in humans(8). Third, given that functional homotopy in humans decreases globally from childhood to middle adulthood(6), we used a longitudinal design to investigate whether the zebra finch brain exhibits a similar trajectory along the developmental process of song acquisition.

## Results and Discussion

### Zebra Finch rs-fMRI Data Demonstrate Comparable Properties to Human Data

Our rs-fMRI data were of comparable quality to rs-fMRI measurements in humans. For each functional scan, the temporal signal-to-noise ratio (*tSNR*), a common metric of rs-fMRI quality(23), was within the range reported for humans(24) (Extended Data Figure 1, panel a). Additionally, as in human data, the zebra finch rs-fMRI signals exhibited power spectrums with a 1/*f* (frequency) distribution, with the greatest power at low frequencies (Extended Data Figure 1, panel b)(25, 26). Furthermore, we noted significant FC among brain regions known to have ipsilateral SC, consistent with known relationships between FC and SC in humans(27) (see Supplementary Materials; Extended Data Figure 1, panel c). For instance, we noted a trend toward positive FC for L HVC – L RA for birds aged P45-90 (*t*(13) = 1.82, *p* = 0.092). Moreover, we noted that the FC of this connection increased significantly with age, from P25 through P90 (*β* = 0.46, *t*(17) = 2.25, *p* = 0.038), consistent with its structural developmental trajectory(28). Additional analyses for known anatomical connections can be found in Supplementary Materials.

### Homotopic Functional Connections are among the Strongest within the Song Network

To test our primary hypothesis, we analyzed homotopic FC across the major telencephalic regions of the song network, including sensory (auditory forebrain (here: caudomedial nidopallium (NCM) and caudomedial mesopallium (CMM)), and Field L), sensorimotor (lateral magnocellular nucleus of the anterior nidopallium, LMAN; Area X; and HVC (proper name), and premotor areas (robust nucleus of the arcopallium, RA). We implemented a linear mixed-effects model predicting ROI-to-ROI FC, controlling for age and subject-specific effects. This revealed that homotopic connections were significantly stronger than both ipsilateral (*B* = 0.16, *t*(1250) = 9.10, *p* < 0.001) and heterotopic (i.e., non-homotopic interhemispheric) connections (*B* = 0.16, *t*(1250) = 9.49, *p* < 0.001). Greater homotopic FC strength was not explained by the Euclidean distance between ROIs, the average volume of ROI pairs, nor the between-subjects effects of physiological parameters, as the FC strength of homotopic connections remained higher than that of other connection types after controlling for these nuisance variables (homotopic vs. ipsilateral: *B* = 0.16, *t*(981) = 9.13, *p* < 0.001; homotopic vs. heterotopic: *B* = 0.11, *t*(981) = 6.22, *p* < 0.001). Notably, the relative strengths of these connection types are consistent with findings in humans and macaques(3).

We next analyzed homotopic FC with respect to individual song network ROIs. We identified significant homotopic FC among all ROIs except for RA (false discovery rate corrected, *pFDR* < 0.05; Figure 2), although RA homotopic FC was significant prior to correction for multiple comparisons (*Z* = 0.06, *pFDR* = 0.055, *p-uncorrected* = 0.011). These analyses indicate that functional homotopy is a global feature of the song network, and is evident within almost all of its individual components, even without direct homotopic SC.

**Fig. 1.**
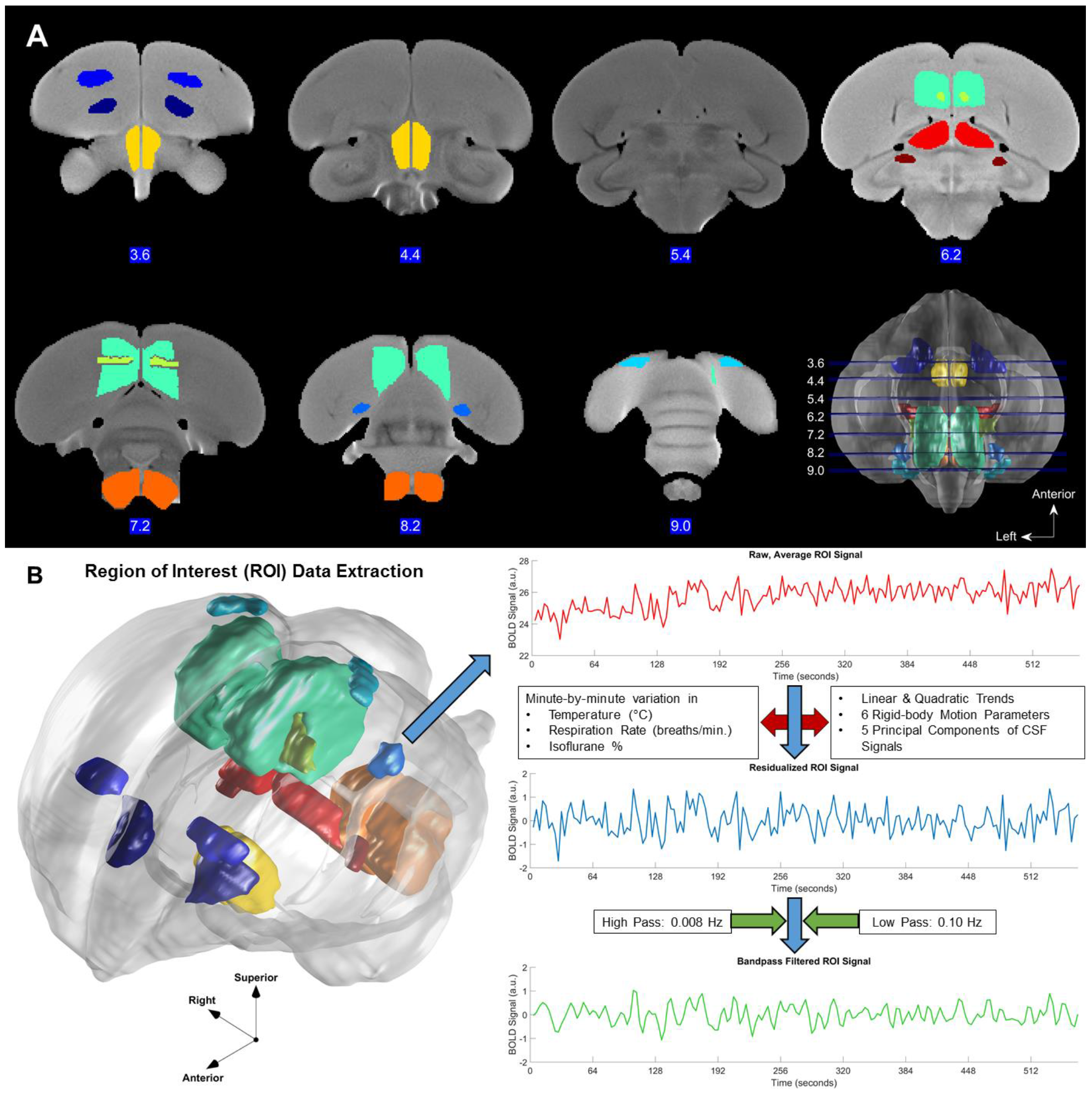
ROI delineation and preprocessing. **A:** Slice mosaic display of the male zebra finch brain and ROIs for the: ***Song Network:*** *Midnight Blue:* Area X; *Royal Blue:* LMAN; *Light Green:* Auditory Forebrain; *Light Yellow:* Field L; *Blue:* RA; *Teal:* HVC; ***Homotopic-SC ROIs:*** *Yellow:* Medial Diencephalon; *Orange:* Medulla; *Red:* Uva; *Maroon:* DM. Brain slices shown are 3.6, 4.4, 5.4, 6.2, 7.2, 8.2, and 9.0 millimeters from the anterior tip of the telencephalon. **B:** *Left:* ROIs displayed within a 3D rendering of our custom structural brain template. 3D axes show brain orientation. *Right:* the basic data extraction and preprocessing steps are shown using data from left RA in male 1 at P25. Signals extracted from all voxels of an ROI are averaged (top), nuisance regression is performed (middle), and ROI signals are bandpass filtered (bottom).

**Fig. 2.**
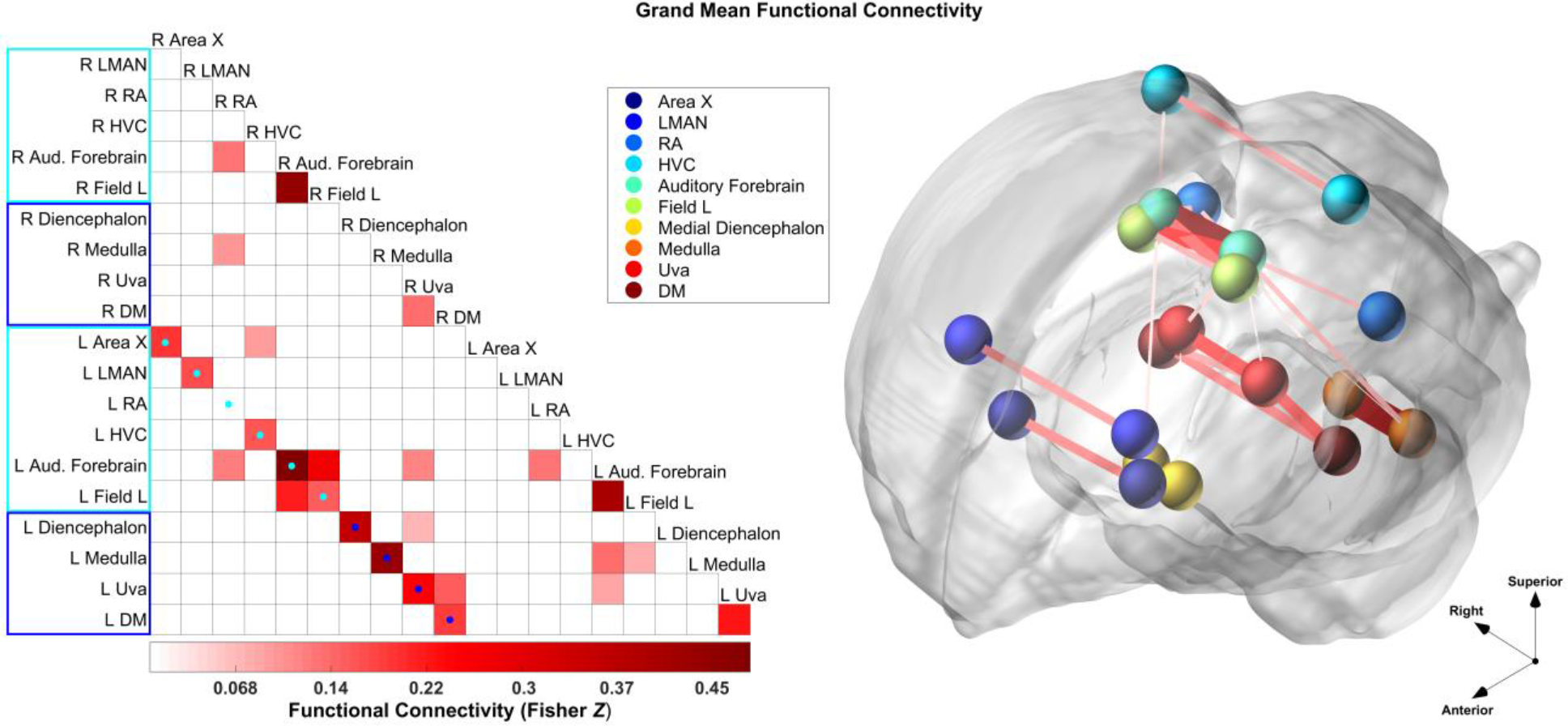
A prominent pattern of homotopic FC. *Left:* Grand mean FC matrix. Homotopic connections are displayed on the diagonal highlighted by teal and blue dots: Area X: *Z* = 0.19; RA: *Z* = 0.06; LMAN: *Z* = 0.17; HVC: *Z* = 0.16; Auditory Forebrain: *Z* = 0.48; Field L: *Z* = 0.15; Diencephalon: *Z* = 0.36; Medulla: *Z* = 0.43; Uva: *Z* = 0.24; DM: *Z* = 0.19. Teal boxes and dots: the song network ROIs (no direct homotopic SC) and homotopic connections, respectively; blue boxes and dots: homotopic-SC ROIs and homotopic connections, respectively. The red intensity scale corresponds to average Fisher *Z*-transformed Pearson product-moment correlations across all males and ages. Significant correlations are displayed using a red color-scale (*pFDR* < 0.05), whereas non-significant correlations, after FDR correction, are white. *Right:* The grand mean FC network overlaid on a 3D-rendering of the group-averaged template brain. Spheres are the centroids of each corresponding ROI listed in the legend. Connections depict the Fisher *Z* values (left panel), with line thickness and color proportional to the association strength. 3D axes show brain orientation.

### Structural Connectivity Predicts Greater Functional Homotopy in Zebra Finches

In humans and macaques, homotopic connections which have direct SC via the CC generally exhibit stronger FC than the minority of homotopic connections which lack direct SC(3). However, no direct homotopic SC projections have been identified between telencephalic zebra finch song network ROIs; thus, these connections can be classified as homotopic non-structurally connected (*homotopic-nSC*)(13). To gauge whether homotopic SC serves to augment homotopic FC in the zebra finch brain, as in the Eutherian brain(3), we added a set of non-telencephalic homotopic ROI pairs that are known to be structurally connected (*homotopic-SC*; medulla, medial diencephalon, nucleus Uvaeformis of the thalamus (Uva), and the dorsomedial nucleus of the intercollicular complex (DM)(13); see Supplementary Materials). As expected, all homotopic-SC ROIs exhibited significant homotopic FC (all *pFDR* < 0.05; Figure 2). A linear mixed-effects model predicting FC across homotopic connections revealed that homotopic-SC connections were significantly stronger than homotopic-nSC connections (*B* = 0.10, *t*(187) = 3.00, *p* = 0.003; Figure 3) even after controlling for nuisance covariates (*B* = 0.08, *t*(142) = 2.38, *p* = 0.019). Thus, although the song system displays substantial homotopic FC without direct SC, homotopy is further strengthened by SC in the zebra finch, as in humans.

**Fig. 3.**
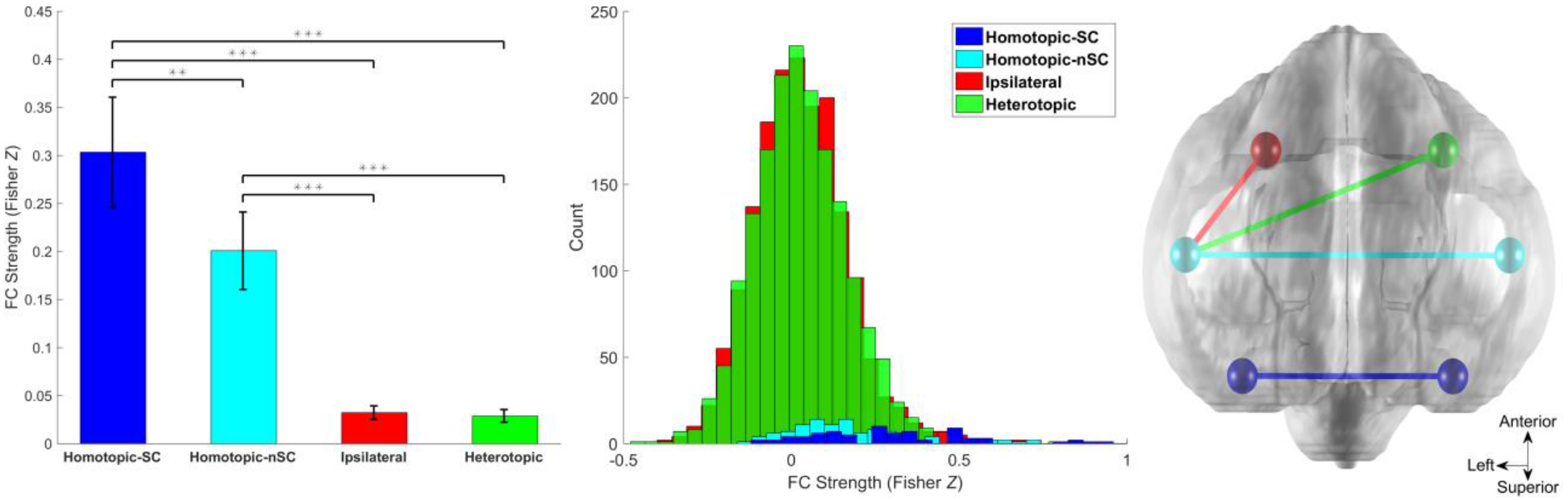
Relative FC of different connection types. *Left:* Bar plot showing mean FC values as Fisher *Z*-transformed Pearson product-moment correlations apportioned by connection type. Error-bars represent 95% confidence intervals. *** = *p* < 0.001, ** = *p* < 0.01. *Middle:* Histograms showing the distribution of FC values for each connection type. *Right:* A schematic illustrating each connection type overlaid on a superior view of the zebra finch brain template. 3D axes show brain orientation.

### Male Zebra Finches Show Less Functional Homotopy than Females, Consistent with More Lateralization, within the Song Network

To test our second hypothesis regarding sex differences in song network homotopic FC, we collected rs-fMRI data from six female zebra finches at P25, P45, and P65 (see Supplementary Materials). Similar to males, homotopic FC was significantly stronger than ipsilateral FC in females (*B* = 0.31, *t*(1137) = 13.21, *p* < 0.001) even when controlling for Euclidean distance and mean ROI volume (*B* = 0.28, *t*(1135) = 12.98, *p* < 0.001). More importantly, we found that homotopic FC, relative to other connection types, was greater in females than males (female*homotopic vs. ipsilateral FC: *B* = 0.10, *t*(4741) = 4.12, *p* < 0.001; female*homotopic vs. heterotopic FC: *B* = 0.10, *t*(4741) = 4.39, *p* < 0.001). Notably, this sex difference was greater for the song network (homotopic-nSC) than for homotopic-SC ROIs (female*homotopic-nSC vs. ipsilateral: *B* = 0.12, *t*(4739) = 3.95, *p* < 0.001; female*homotopic-nSC vs. heterotopic: *B* = −0.12, *t*(4739) = −4.16, *p* < 0.001; female*homotopic-SC vs. ipsilateral: *B* = 0.07, *t*(4739) = 1.84, *p* = 0.066; female*homotopic-SC vs. heterotopic: *B* = −0.07, *t*(4739) = −2.01, *p* = 0.044), suggesting possible functional specificity. Prior findings in humans^14^ and zebra finches(12) indicate that brain networks for vocal learning may be lateralized across phylogeny. Given that only male zebra finches learn to sing, it therefore follows that male zebra finches would show reduced functional homotopy and increased lateralization in the song network compared to females, who cannot sing.

### Homotopic FC Decreases with Age in Male Zebra Finches

In humans, functional homotopy decreases globally from early childhood through middle adulthood(6). To test our third hypothesis, that a similar pattern of increasing lateralization occurs over the developmental time-course of song learning, we implemented a linear mixed-effects model predicting ROI-to-ROI FC via age, connection type, and their two-way interaction using the male zebra finch longitudinal rs-fMRI data. Consistent with human findings, zebra finch homotopic FC decreased significantly more with age than did ipsilateral FC (*β* = −0.15, *t*(3604) = −2.03, *p* = 0.042), whereas heterotopic FC did not differ developmentally from ipsilateral FC (*β* = 0.03, *t*(3604) = 0.99, *p* > 0.30). The behavioral relevance of developmental increases in lateralization, whether for human cognition or zebra finch song learning, merits further study. The cross-species parallel noted here indicates that the zebra finch might serve as an important model organism for understanding this developmental phenomenon.

### Voxel-wise Analyses Demonstrate a Medially-biased, Brain-wide Pattern of Functional Homotopy

To further interrogate the distribution of homotopic FC agnostically, without pre-defining ROIs, we implemented the experimenter-blind voxel-mirrored homotopic connectivity (VMHC)(6) method. We created a symmetrical male brain template and normalized all rs-fMRI images to this template for this analysis. VMHC revealed a pattern of strong medial FC strikingly similar to the voxel-wise pattern observed in the human brain(4, 6): homotopic FC was higher within medial portions of the zebra finch brain, spanning the medulla to the superior telencephalon, with reduced homotopic FC in more lateral regions (Figure 4). Notably, the midline homotopic FC was not an artifact of spatial smoothing, as we discarded data from a four-voxel radius of the midline within each hemisphere. Six significant clusters of homotopic FC were identified within the VMHC voxel-wise pattern (all *p* < 0.05, family-wise error corrected; Figure 4, Extended Data Table 2). The proportion of voxels from significant clusters that overlapped with song network ROIs (8.2%) greatly exceeded that which would be predicted by chance (permutation test *p* < 0.0001). These VMHC results strongly confirm that zebra finch functional brain networks are characterized by widespread functional homotopy. Additionally, the fact that our experimenter-drawn song network ROIs exhibited more prevalent homotopic FC than a random sampling of brain voxels suggests that bilateral coordination may be of particular functional importance for song, a complex and distributed multimodal brain process.

**Fig. 4.**
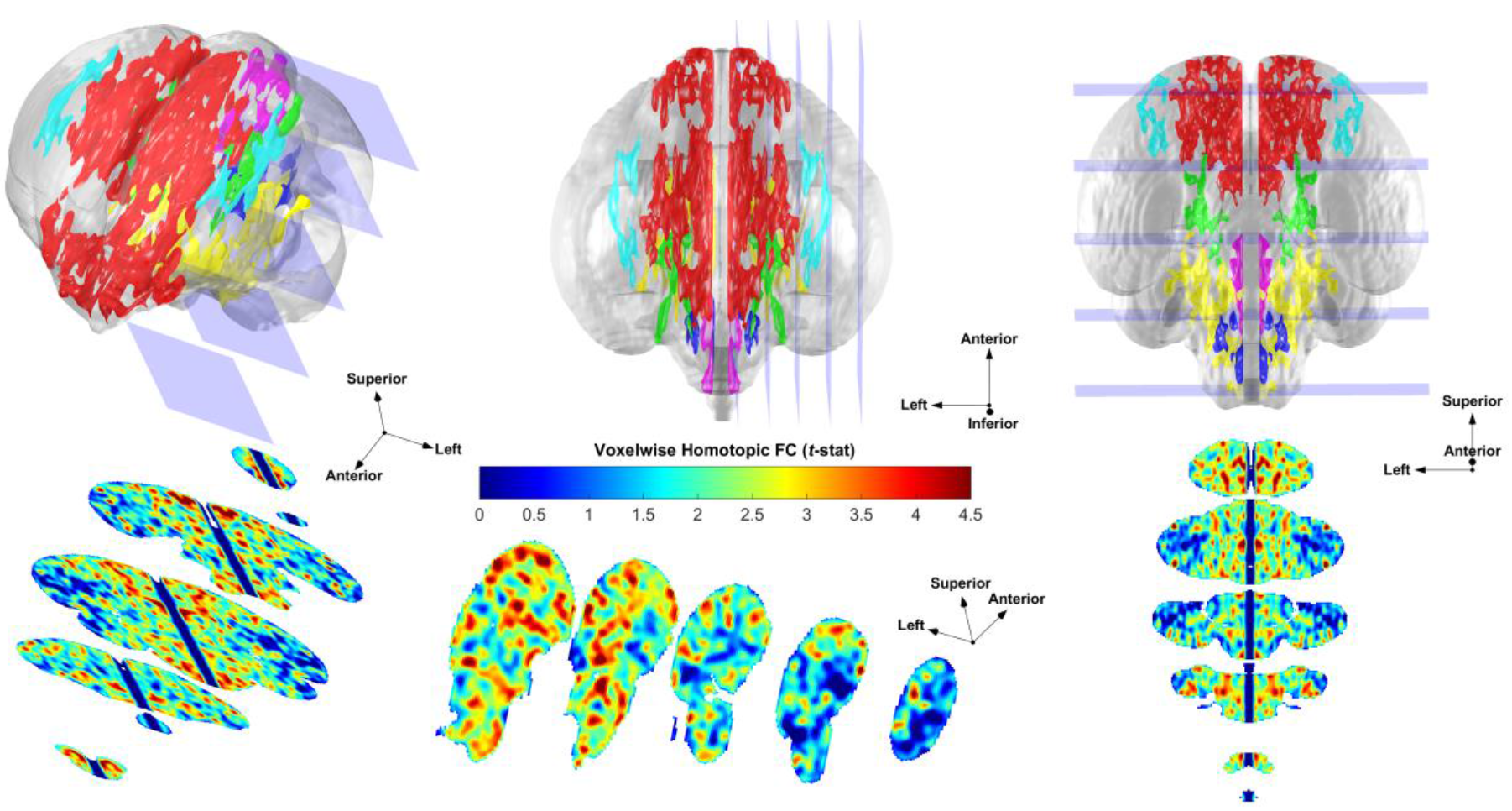
Voxel-wise homotopic resting-state FC. *Left Column*: ventral view and coronal slices. *Middle Column*: superior view and sagittal slices. *Right Column*: posterior view and axial slices. *Top row:* Six significant clusters were identified: cluster 1 (magenta) was located within the cerebellum; cluster 2 (green) overlapped Uva and parts of Auditory Forebrain, Field L, and RA; cluster 3 (blue) overlapped Medulla; cluster 4 (red) overlapped portions of Auditory Forebrain, LMAN, Area X, Field L, and Uva; cluster 5 (cyan) overlapped unlabeled regions of superior and lateral telencephalon; and cluster 6 (yellow) overlapped DM, Medial Diencephalon, and Uva (see Extended Data Table 2). Significant homotopic FC clusters overlapped the following proportions of experimenter-drawn ROIs: DM (44.7%), LMAN (15.1%), Uva (9.3%), Auditory Forebrain (9.0%), Medulla (6.6%), Field L (6.6%), Medial Diencephalon (3.4%), RA (2.9%), and Area X (2.4%).

### Conclusion

Our results demonstrate widespread functional homotopy in an organism that lacks a CC in the course of normal ontogeny. This indicates that functional homotopy may predate the CC, supporting interhemispheric coordination more broadly across phylogeny. Accordingly, rather than serve as the direct cause of functional homotopy, the CC may have evolved to provide finer control over bilateral integration and segregation. Consistent with this notion, it has recently been shown that precise homotopic wiring of the CC in the postnatal rodent brain requires preexisting functional homotopy(29). It is therefore plausible that ancestral pathways such as the anterior commissure(30) or symmetrical ascending bilateral inputs(13) are sufficient to provide the balance of interhemispheric integration and specialization required for complex multimodal cognition. The ability to conduct longitudinal investigations during a short sensitive period for vocal development makes the zebra finch an ideal model organism for investigating the behavioral relevance of this balance, allowing experimental manipulations not possible in humans.

## Materials and Methods

### Overview

We collected rs-fMRI scans for male zebra finches at four Posthatch (P) ages to capture the beginning, middle, and end of the song learning process (P25, P45, P65, and P90; n=5 each age). As previously validated in task-based fMRI, the birds were lightly anesthetized with isoflurane (0.5-2%) for the duration of scanning(31). Following implementation of a custom preprocessing pipeline to analyze small brains across development, we calculated FC as the Fisher’s *Z*-transformed Pearson correlation between rs-fMRI signals extracted from manually delineated ROIs encompassing major components of the song system (Figure 1, Extended Data Table 1). This FC measure reliably detects functional homotopy in Eutherians(3, 4, 6) and provides results consistent with alternative measures of brain connectivity(1, 2).

### Animals

All procedures were approved by the Institutional Animal Care and Use Committee of the University of Chicago. Five male zebra finches were scanned longitudinally on days P25, P45, P65 and P90. A P90 MRI scan was not obtained for one male, yielding a final sample of 19 functional series. Due to scanner availability, not all birds could be scanned on the exact target days: P25 (range: P24-P26), P45 (range: P44-P46), P65 (range: P64-P67), and P90 (range: P88-P91). All birds were maintained under a 14/10 hour light/dark photoperiod throughout the experiment, with food provided *ad libidum*.

### Scanning Procedure

MRI data collection was conducted at the MRIS Facility of the University of Chicago. Upon arrival, zebra finches were anesthetized using an admixture of oxygen and isoflurane gas (1.5-2.25%). Subsequently, a light maintenance dosage of isoflurane was maintained throughout the experiment (0.5-2%), and was administered via a tube fitted to the zebra finch beak. The magnitude of hemodynamic responses are reduced by isoflurane anesthesia, but the shape of hemodynamic responses remains similar to awake animals(32). The use of isoflurane is well-established in task-based fMRI studies of zebra finches(31, 33). Following anesthesia administration, birds were fitted with a temperature probe and respiration monitoring pad, allowing for body temperature, respiratory rate, and isoflurane percentage to be recorded at one-minute intervals throughout the scanning period. The zebra finches were wrapped in a felt cloth to help maintain body temperature. Upon insertion into the scanner, a warm air feedback system was used to maintain body temperature within a normal physiological range (40.0 ± 0.2 °C).

### Imaging Data Acquisition

Neuroimaging data were acquired using a 30 cm bore 9.4 Tesla Bruker small animal MRI scanner. A TurboRARE-T2 Multislice anatomical scan was acquired first during each scanning session (TR = 3.5 s, TE = 20 ms, Matrix Size: 256 × 256, in-plane resolution = 70.3 μm × 70.3 μm, slice-thickness = 200 μm, 59 slices, 9 averages). Resting-state spin-echo T2*-weighted MR images were then acquired (TR = 1.07 s, TE = 26.97 ms, RARE Factor = 12 (3 repetitions per acquisition), Matrix Size: 128 × 36, in-plane resolution = 141 μm × 500 μm, slice-thickness = 750 μm, 15 slices). Slices were acquired in an interleaved, ascending order. 180 volumes were acquired consecutively with an effective sampling rate of 3.20 seconds and a total resting-state scan time of 9.60 minutes. To avoid T1-equilibration effects, the first five volumes of each functional series were discarded. A spin-echo pulse sequence was utilized due to observations that gradient-echo imaging may be particularly vulnerable to susceptibility artifacts in zebra finch whole-brain fMRI(34).

### Preprocessing

Image preprocessing was completed using a combination of Advanced Normalization Tools (ANTs)(35, 36), Statistical Parametric Mapping (SPM12)(37), and custom Matlab scripts. First, DICOM files were converted to NIfTI format using our custom MRIqual toolbox (E.A.L. & M.G.B., 2017). Second, the “N4” bias field correction algorithm(38) was used to correct each anatomical volume for magnetic field intensity inhomogeneity. Third, an average of the bias-corrected anatomical scans was selected as an unbiased starting point for symmetric group-wise normalization (SyGN)(39) and template-building using ANTs. This process involves iteratively registering each anatomical scan to a custom brain template using affine and nonlinear transformations. As image registration improves, the custom template is recursively updated at each iteration. Cross-correlation was implemented as the similarity metric for image comparison, and four template construction iterations were utilized.

Fourth, functional scans were corrected for slice timing differences using SPM. Slice timing correction may be particularly essential in the case of longer duration TRs(40), and it is recommended that this step be performed prior to realignment for interleaved slice acquisitions(41). Fifth, each 3D functional volume was realigned to the first volume in its series using ANTs, and the six rigid-body motion parameters were retained from each transformation applied. Sixth, the first functional volume of each series was coregistered to the corresponding bias-corrected anatomical volume via affine transformation in ANTs. To minimize the number of interpolations applied, all affine and nonlinear transformations obtained from functional realignment, coregistration, and structural normalization were combined into 3D volume-specific deformation fields using the ANTs routine “antsApplyTransforms.” Seventh, deformation fields were applied in a single step to each slice timing corrected 3D functional volume. Lastly, we assessed the quality and consistency of spatial normalization by computing voxel-wise Pearson correlations between the custom template and normalized anatomical scans across voxels (*r_mean_* = 0.94, *SD* = 0.01, *r_min_* = 0.91), as well as between the normalized functional scans averaged across time and the custom template (*r_mean_* = 0.87, *SD* = 0.02, *r_min_* = 0.81), with both analyses indicating a robust and consistent spatial normalization across scans.

### Region of interest (ROI) delineation

Using our NeuroViz Matlab toolbox (E.A.L. & M.G.B., 2017), coauthor S. London manually delineated six key components of the zebra finch song system within each hemisphere on our anatomical brain template (homotopic non-structurally connected, *homotopic-nSC* ROIs). Additionally, four regions that included brain areas known to receive bilateral inputs or direct homotopic structural connectivity were delineated: the medulla, medial diencephalon, thalamic nucleus uvaeformis (Uva), and the dorsomedial subdivision of the intercollicular nucleus (DM)(13, 42–44). These latter ROIs were considered homotopic structurally connected ROIs (*homotopic-SC*). For further details on the homotopic structural connectivity of these regions, see Supplementary Materials. This yielded a total of 20 ROIs, comprised of 10 bilateral homotopic pairs (see Figure 1A, Extended Data Table 1).

### Data extraction and denoising

For each functional series, resting-state BOLD time series were extracted and averaged across the voxels of each ROI. We implemented denoising procedures using MRIqual and custom Matlab scripts. Specifically, a nuisance regression was performed in which the six rigid body motion parameters(45), the first five principal components of cerebrospinal fluid (CSF) signals, respiratory rate, body temperature, isoflurane percentage, and linear and quadratic trends(46) were removed from each ROI time series. Physiological variables recorded at one-minute intervals were interpolated using a cubic spline function to the temporal resolution of our functional series for nuisance regression. Respiratory rate data were not usable for four of the 19 functional series scans (Male 1 P25, Male 2 P60, Male 3 P90, & Male 4 P90) because peaks in the respiratory signal were frequently (but non-uniformly) double-counted by the respiration monitoring equipment, yielding invalid data. Following nuisance regression(47), the residualized ROI time series were bandpass filtered (range: 0.008 to 0.1 Hz). See Figure 1B for a visual depiction of this process.

### rs-fMRI validation analyses

Temporal Signal-to-Noise Ratio (*tSNR*)(23) was calculated across all brain voxels and averaged for each resting-state functional series using MRIqual. *tSNR* was calculated prior to preprocessing to avoid removing signal means, which would invalidate *tSNR* calculations. *tSNR* values obtained were compared across scans to assure consistency of image quality, and these were also compared to values cited for human fMRI studies(24). Given the small volume of our functional voxels (~0.05 mm^3^), *tSNR* and similar metrics were expected to be markedly lower than in human resting-state scans collected at lower spatial resolution (i.e, more coarse), because *tSNR* is known to positively correlate with voxel volume where larger volume yields higher *tSNR*(23, 48). Notably, the quantification of image quality metrics such as these may be particularly invaluable for novel scanning paradigms and/or for use with model organisms, for which scanning parameters are often less well-established(49).

Power spectrums for each resting-state functional series were calculated within the approximate human resting-state frequency range (~0.008-0.1 Hz)(50) after the completion of preprocessing. Average power was calculated via Welch’s power spectral density estimate, implemented using the Matlab function pwelch.m and integrated within MRIqual. The power spectrums obtained were compared to those reported in the human resting state literature(51, 52) and in the murine literature, which routinely utilizes isoflurane anesthesia(25, 26). Both literatures have consistently identified 1/*f* power spectrums, with resting-state power highest in low frequency bands (< 0.1 Hz). In particular, resting-state time series obtained from isoflurane anesthetized mice(25) and rats(26) have demonstrated peaks in the power spectrum at very low frequencies (~ 0.01 Hz).

Anatomical connectivity is known to moderately predict functional connectivity at rest in humans(27). However, distinct brain networks are associated with the resting-state as compared to various task states(53). Therefore, it is uncertain to what extent we might expect to observe FC between previously delineated premotor and motor pathways during the resting-state. However, we proceeded to investigate whether there was consistency between the FC measured in the present study and structural connectivity known from prior research. In particular, an ipsilateral projection from HVC to RA comprises an important component of the descending motor pathway for song production, and this anatomical connection has been observed previously using diffusion tensor imaging (DTI) in starlings(54). Notably, projections from HVC to RA do not project into RA until after ~P30, and they are thought to increase in density and efficacy from P30 to adulthood (>P90)(28). Therefore, we restricted our search for any analogous FC between these regions to birds P45 or older, and we additionally investigated whether FC was predicted by age. Additional investigations of known ipsilateral projections are also detailed (see Supplementary Materials, Extended Data Figure 1, panel c).

### Homotopic FC analyses

ROI-to-ROI FC was computed separately for each functional time series. FC was defined as the Fisher’s *Z*-transformed Pearson correlation between the preprocessed time series of a pair of ROIs. To identify the statistically significant features of the zebra finch brain network across all birds, one-sample *t*-tests were conducted to determine whether Fisher *Z* values for a given ROI pair significantly differed from zero across functional series. Then, a grand mean ROI-to-ROI FC network was constructed by averaging Fisher’s *Z* values across all functional series (i.e., all birds and time points). To account for multiple comparisons, functional connections of the grand mean network were thresholded at *p-FDR* < 0.05.

We next sought to determine whether homotopic connections exhibit increased FC strength within the zebra finch brain, as has been shown consistently in mammals. To investigate these questions, a linear mixed-effects model was implemented in which ROI-to-ROI FC across all ROIs served as the criterion variable. Connection type (homotopic, ipsilateral, and heterotopic) was entered as a three-level nominal categorical variable, with ipsilateral coded as the reference category. We also controlled for age and included a random intercept for bird. This linear mixed-effects model was optimized for maximum likelihood using the Matlab function “fitlme.” A contrast was performed for the effect of homotopic vs. ipsilateral connections.

### Additional controls

We sought to control for a variety of nuisance factors known to influence resting-state FC. For instance, the Euclidean distance between a pair of ROIs is known to be inversely associated with the FC strength of the pair(27, 55, 56). Additionally, increased ROI volume is associated with increased FC strength(57, 58). Similarly, concerns have been raised regarding the potential for regional *tSNR* variations to influence FC measures(59). Therefore, we added these three covariates to our previously described mixed-effects model.

Additionally, although our preprocessing sequence included control of within-scan fluctuations in body temperature, respiration rate, and isoflurane percentage, these controls did not account for possible *between-scan* or *between-bird* variance in ROI-to-ROI FC explained by these factors (e.g., if average isoflurane percentage over the duration of a functional acquisition negatively predicts global FC strength(60)). Thus, we added average body temperature, average respiration rate, and average isoflurane percentage over the duration of functional acquisition as additional between-subjects covariates in our mixed-effects model.

### Homotopic-SC vs. homotopic-nSC

We sought to investigate whether FC was significantly stronger between homotopic-SC ROI pairs than between homotopic-nSC ROI pairs. Therefore, we implemented a linear mixed-effects model, here considering only the FC of homotopic connections. Connection type was added as a binary categorical variable (homotopic-SC vs. homotopic-nSC), and age and a random intercept for bird were included as covariates. In a follow-up regression, we also included the covariates described in “Additional controls.”

### Homotopy as a function of sex and development

Male and female zebra finches exhibit a number of structural and functional brain dimorphisms, and song production occurs only in males(61). Therefore, we sought to investigate whether the strength of homotopic FC differed between sexes. We utilized a previously collected pilot sample consisting of six females, two scanned at each of P25, P45, and P65. See the Supplementary Materials for details on image acquisition and preprocessing for this sample. First, to gauge whether females also exhibited homotopic FC, we recomputed the mixed-effects model described in “Homotopic FC analyses” using only female data. Secondly, we concatenated all male and female data and re-implemented this mixed-effects model, additionally including the main effect of sex and an interaction term between connection type and sex. Finally, we parsed connection type into a four-level categorical variable (homotopic-SC, homotopic-nSC, ipsilateral, heterotopic), allowing us to gauge whether the sex by connection type interaction was specific to either the homotopic-SC or homotopic-nSC connection type.

In human subjects, global homotopic FC has been shown to decrease from early childhood through middle age (and later reversing), a trend which may reflect the development of increasing lateral specialization(6). Thus, we sought to investigate whether the FC strength of homotopic connections demonstrated a similar trajectory in male zebra finches. We therefore added a term for the interaction between age and connection type to the model described above in “Homotopic FC analyses.” We then conducted a follow-up analysis parsing the connection type variable into four levels (homotopic-SC, homotopic-nSC, ipsilateral, heterotopic), allowing us to gauge age effects specific to each connection type.

### Voxel-wise distribution of homotopic FC

We sought to investigate the voxel-wise distribution of homotopic FC across the entire brain. By illuminating regional clusters of homotopic FC, such an analysis has the potential to be informative for future studies investigating putative neural pathways underlying interhemispheric coordination in the avian brain. To accomplish this, we conducted a voxel-mirrored homotopic connectivity (VMHC)(6) analysis, which quantifies FC between each voxel in one hemisphere with its mirrored counterpart in the contralateral hemisphere. The VMHC analysis proceeded as follows: first a symmetrical group-wise brain template was formed by mirroring the right hemisphere of our custom brain template across the midline. Each individual anatomical scan was then renormalized to this symmetrical brain template. The transformation parameters obtained from normalization were then applied to each corresponding functional series, using the same procedure as described previously. Subsequently, each functional scan was smoothed using a Gaussian kernel (281.2 × 281.2 × 800 μm FWHM, corresponding to 4 × 4 × 4 voxels in anatomical dimensions and 2 × 0.56 × 1.07 voxels in functional dimensions) to help account for any registration errors and/or individual-specific brain anatomy. Next, Fisher’s *Z*-transformed Pearson correlations were computed between each pair of homotopic brain voxels, and voxel-wise *t*-statistics were calculated to summarize the strength of homotopic FC across birds and time points. Finally, all voxels within a distance of 281.2 μm (i.e., four voxels) from either side of the midline were set to zero to avoid obtaining artefactual homotopic FC as a result of spatial smoothing (for FWHM = 281.2 μm, σ ≅ 119.4 μm, yielding an approximate cut-off for signal contamination at 3 * σ = 358.2 μm, or 5.10 voxels).

Significant clusters of homotopic FC were detected using the cluster-extent based thresholding method (primary or voxel-level threshold: *p* < 0.001, cluster-extent threshold: *p-FWE* < 0.05)(62). Overlap between significant clusters of homotopic FC with our previously delineated homotopic-nSC song control nuclei and homotopic-SC nuclei was calculated as the number of intersecting voxels divided by the total number of voxels encompassed by a given nuclei or group of nuclei. A two-tailed *Z*-test for proportions was used to contrast the proportion of overlap for homotopic-nSC ROIs with the proportion of overlap for homotopic-SC ROIs. Finally, to compare the amount of overlap between significant homotopic FC clusters with either homotopic-nSC or homotopic-SC ROIs versus chance, we utilized a permutation test in which the indices of significant cluster voxels were randomly permuted 10,000 times, and the proportion of overlap with our previously delineated ROIs was calculated in each case. The original observed overlap proportion was then compared to the null distribution of overlap proportion obtained via permutations to derive a non-parametric *p*-value.

### Figure Creation

Figures 1–4 were created using our NeuroViz Toolbox and custom Matlab scripts.

## Data availability

The neuroimaging data that support the findings of this study are available from the corresponding author upon reasonable request.

## Acknowledgements

This research was supported in part by a grant from the Big Ideas Generator (BIG) at the University of Chicago to S.E.L. and M.G.B and by a grant from the National Science Foundation (NSF) BCS-1632445 to M.G.B. We thank the staff of the MRIS facility and Frank Ibarra for their technical assistance, and Elisa Gores for bird care.

## Author Contributions

The study was conceived by M.G.B., S.E.L., and E.A.L. Neuroimaging data were collected by all authors. Neuroimaging data preprocessing and statistical analyses were conducted by E.A.L. E.A.L., M.G.B., K.E.S., and S.E.L. wrote the manuscript with contributions from all authors.

